# Co-variation of viral recombination with single nucleotide variants during virus evolution revealed by CoVaMa

**DOI:** 10.1101/2021.09.14.460373

**Authors:** Shiyi Wang, Stephanea L. Sotcheff, Christian M. Gallardo, Elizabeth Jaworski, Bruce E. Torbett, Andrew L. Routh

## Abstract

Adaptation of viruses to their environments occurs through the acquisition of both novel Single-Nucleotide Variants (SNV) and recombination events including insertions, deletions, and duplications. The co-occurrence of SNVs in individual viral genomes during their evolution has been well-described. However, unlike covariation of SNVs, studying the correlation between recombination events with each other or with SNVs has been hampered by their inherent genetic complexity and a lack of bioinformatic tools. Here, we expanded our previously reported CoVaMa pipeline (v0.1) to measure linkage disequilibrium between recombination events and SNVs within both short-read and long-read sequencing datasets. We demonstrate this approach using long-read nanopore sequencing data acquired from Flock House virus (FHV) serially passaged *in vitro*. We found SNVs that were either correlated or anti-correlated with large genomic deletions generated by nonhomologous recombination that give rise to Defective-RNAs. We also analyzed NGS data from longitudinal HIV samples derived from a patient undergoing antiretroviral therapy who proceeded to virological failure. We found correlations between insertions in the p6^Gag^ and mutations in Gag cleavage sites. This report confirms previous findings and provides insights on novel associations between SNVs and specific recombination events within the viral genome and their role in viral evolution.

## Introduction

Recombination within viruses, and particularly RNA viruses, is a powerful driving force behind their evolution and adaptation (1). Recombination may result in manifold outcomes, including the reshuffling of advantageous or deleterious mutations among homologous viral genomes, insertions or duplications of host or viral genomic segments, or deletion of genomic segments to generate Defective Viral Genomes (DVGs). These genetic changes can have a dramatic impact upon viral evolution, fitness, viral intra-host diversity, and the development of resistance to antivirals (2–5). Due to the dense and compact nature of viral genomes, adaptive mutations rarely occur in isolation but rather mutations are often correlated with one another (6–8). Characterizing how or whether individual adaptions are correlated is essential to understand the mechanisms whereby viral strains emerge and adapt to their environment, for example, in response to anti-viral therapy or immune pressure.

A number of approaches have been documented to characterize the emergence of correlated viral adaptations (9). Classical approaches use consensus-level viral genomic data derived from multiple individuals/hosts deposited in curated databases such as GISAID, Stanford HIVdb, and others (10–12). More recently, approaches have focused on extracting evidence for mutational covariation in Next-Generation Sequencing (NGS) data that reports on virus intra-host genetic diversity. Broadly, these methods either directly assess the frequency of detected mutations found co-occurring within NGS reads, or infer correlated mutations using probabilistic or mathematical models (6, 13–17). However, there are no current approaches aimed at determining the correlation of recombination events, such as insertions, deletions, or duplications, with each other or with Single Nucleotide Variants (SNV). Characterizing these correlations may be necessary for understanding why certain recombination events might be selected for and their role in viral evolution.

We recently reported a method that measures linkage disequilibrium of SNVs within viral NGS datasets (16). This approach, called CoVaMa (<u>Co</u>-<u>Va</u>riation <u>Ma</u>pper), built a large matrix of contingency tables corresponding to every possible pair-wise interaction of correlated nucleotides within a viral genome. Each contingency table was a 4×4 matrix with columns and rows corresponding to the bases ‘ATGC’ for each nucleotide position and was populated by extracting every possible pair of co-mapping nucleotides within individual reads. From each 4×4 table, all possible 2×2 tables were extracted to measure linkage disequilibrium (LD), thus reporting evidence of mutational covariation. This basic process is a non-heuristic and brute-force approach implemented in a series of python scripts. We demonstrated the utility of this approach by identifying correlated mutations within the genomic RNAs of Flock House virus (FHV) and within anti-viral inhibitor-resistant HIV patient samples.

However, this approach retained many limitations, including the requirement for a gapless alignment and the inability to consider the correlation of other types of mutations besides SNVs. As we and others have shown (4, 18–20), recombination events and InDels comprise an important component of the viral intra-host diversity. However, while there is evidence of the co-evolution of SNVs and other types of mutations, high-throughput tools to assess the epistatic linkage between recombination events and/or SNVs remain lacking. The requirement for a gapless alignment also prevented the analysis of long-error prone read data, such as nanopore or PacBio data. These data types have the distinct advantage in that they can resolve epistatic associations over much greater distances than Illumina short-read data and can even resolve full-length viral genomes direct from RNA.

Here, we expanded the capabilities of CoVaMa to measure linkage disequilibrium between recombination events found within deep-sequencing reads and SNVs. Recombination events, such as insertions and deletions, can be extracted from gapped-alignments using *HISAT2* (21) and *Bowtie2* (22), or using other tools designed for mapping recombination in virus genomes such as *ViReMa* (23). Moreover, our improved algorithm can accept long-reads generated from Nanopore sequencing platforms. This approach used a similar principle employed by the original version of CoVaMa by generating large matrices of contingency tables corresponding to every pair-wise interaction of each nucleotide position with each detected recombination event in the NGS dataset. We developed a set of criteria that determined when a sequence read excluded the possibility of mapping across a recombination event. This allowed the populating of 4×2 contingency tables for each nucleotide (ATGC) at each genomic position with either the confirmed absence or presence of a recombination event from which the degree of covariation or Linkage Disequilibrium value was calculated.

We demonstrate the utility of this approach by re-analyzing long-read nanopore data that we generated in a recent longitudinal study of Flock House virus (FHV) evolution in cell-culture (24) and short-read Illumina data from a study of clinical HIV patient samples collected during the development of antiretroviral therapy resistance. In the FHV dataset, we identified multiple SNVs that were either correlated or anti-correlated with large deletion events that constituted Defective-RNAs (D-RNAs). We hypothesize that these mutations were adaptive mutations that either allowed the heightened replication of D-RNAs, or that allowed the wild-type full-length genome to escape attenuation by D-RNAs. From the HIV samples, we identified insertions in the PTAP region of the p6^Gag^ that correlated with mutations found in Gag cleavage sites. The location of these linked SNVs and InDels proximal to Gag cleavage sites suggested a role for these adaptions to support drug-resistance development. Overall, CoVaMa provides a powerful tool to characterize the molecular details of viral adaptation and the impact of RNA recombination upon virus evolution. The CoVaMa (v0.7) script is publicly available at https://sourceforge.net/projects/covama/.

## Methods

### Long-read data from serial passaged FHV using CoVaMa

We previously described a serial passaging experiment of Flock House virus (FHV) followed by the analysis of viral genetic changes by the parallel use of short-read Illumina and long-read nanopore sequencing (24). Briefly, pMT vectors containing cDNA of each FHV genomic RNA were transfected into S2 cell in culture and expression of genomic RNA was induced with copper sulphate. After 3 days, 10 ml of supernatant was retained. 1ml of supernatant was passaged directly onto fresh S2 cells in culture, and FHV virions were purified from the remaining supernatant using sucrose cushions and sucrose gradients as described. This process was repeated for a total of nine passages. Encapsidated RNA was extracted from each stock of purified virions and reverse transcribed using sequence-specific primers cognate to the 3’ ends of FHV RNA1 and FHV RNA2. Full-length cDNAs were then PCR amplified in 19 cycles with NEB Phusion using primers targeting the 5’ and 3’ end of each genomic segment. Final PCR amplicons were used as input for Oxford Nanopore Amplicon sequencing protocol by ligating on native barcodes and then the ONT adaptor. Pooled libraries were sequenced on an ONT MinION MkIB with R9 (2017) flowcells. Data was demultiplexed and basecalled using the Metrichor software and using *poretools* (25). Datasets are publicly available in NCBI SRA under the accession code SRP094723. Basecalled data were mapped to the FHV Genomic RNAs (FHV RNA1: NC_004146, 3107 nts) and (FHV RNA2: NC_00414, 1400 nts) using *BBMap*. SAM files of the aligned data were passed to the CoVaMa (Ver 0.7) pipeline using the command lines shown in **Box 1**.

#### Box 1.

**Command lines used to analyze nanopore sequencing reads acquired from each passage of FHV.**

1. Making matrices containing contingency table for each association using CoVaMa_Make_Matrices script: Output_Tag, the name for the output pickle file [Output_Tag].Total_Matrices.py.pi. Data_directory, the folder contains the FHV reference sequence in FASTA file format and the aligned sequences in SAM file format. Contingency tables populated by over 100 reads and the mutant frequency higher than 0.05 were generated and passed to the output pickle file. *python2 CoVaMa_Make_Matrices*.*py [Output_Tag] Data_directory/FHV_Genome_corrected*.*txt--SAM1 Data_directory/FHV_mapping*.*sam --PileUp_Fraction 0*.*05 --Min_Fusion_Coverage 10 NT*
2. Calculating linkage disequilibrium values for each contingency table using CoVaMa_Analyse_Matrices script: [Output_Tag].Total_Matrices.py.pi generated by CoVaMa_Make_Matrices script was used as the input. Linkage disequilibrium information was stored in the output file in TXT file format. A minimum coverage of over 10 pairs of associated nucleotides and recombination events was required for the contingency table to be analyzed. The linkage disequilibrium values (LD) and R square values for each association were normalized by the number of reads populating that contingency table. *python2 CoVaMa_Analyse_Matrices*.*py [Output_Tag]*.*Total_Matrices*.*py*.*pi CoVaMa_output*.*txt --Min_Coverage 10 –-Min_Fusion_Coverage 10 -OutArray -Weighted NT*

### Short-read data from longitudinal HIV patient samples using CoVaMa

We previously described an RT-PCR approach to amplify the *gag-pol* regions of HIV from 93 clinical specimens for Next-Generation Sequencing (6, 26). Briefly, two overlapping cDNA amplicons were RT-PCR amplified using two pairs of primers targeting *gag-pol* (F1: NT 754, R1: 1736; F2: 1569, R2: 2589). Full-length cDNA amplicons were sheared to approximately 175bp and submitted for paired-end sequencing (2×150bp) on an Illumina HiSeq. We chose five longitudinal samples derived from a single patient over six years. Paired-end reads were merged using *BBMerge* and mapped to the HIV genome (HXB2: K03455.1, 9719bp) using ViReMa v0.21 (23) with default settings plus the following parameters (*--X 3 --MicroInDel 5 –BackSplice_limit 25*). As the cDNA amplicon strategy for Illumina sequencing was not directional, we only used the reads mapping to the positive sense viral genome for further analysis. These SAM files were passed to CoVaMa (Ver 0.7) as described in **Box 2**.

#### Box 2.

**Command lines used to analyze NGS sequencing reads acquired from each longitudinal HIV sample.**

1. Making matrices containing contingency table for each association using CoVaMa_Make_Matrices script: Output_Tag, the name for the output pickle file [Output_Tag].Total_Matrices.py.pi. Data_directory, the folder contains the HIV reference sequence for each sample in FASTA file format and the aligned sequences in SAM file format. Contingency tables populated by over 100 reads and the mutant frequency higher than 0.01 were generated and passed to the output pickle file.
  a. For the 15nt insertion event at nt 2158, 15 nucleotides were used to exclude negative recombination event. The analysis region started at nt 1958 and ended at nt 2358.
  b. For the 6nt insertion event at nt 1149, 6 nucleotides were used to exclude negative recombination event. The analysis region started at nt 949 and ended at nt 1349.
  *python2 CoVaMa_Make_Matrices*.*py [Output_Tag] Data_directory/HIV_reference_seq*.*txt –Mode2 Recs --SAM1 Data_directory/HIV_mapping*.*sam --PileUp_Fraction 0*.*01 --Rec_Exclusion 15 -- NtStart 1958 --NtFinish 2358 NT*
2. Calculating linkage disequilibrium values for each contingency table using CoVaMa_Analyse_Matrices script: [Output_Tag].Total_Matrices.py.pi generated by CoVaMa_Make_Matrices script was used as the input. Linkage disequilibrium information was stored in the output file in TXT file format. A minimum coverage of over 100 pairs of associated nucleotides and recombination events was required for the contingency table to be analyzed. The linkage disequilibrium values (LD) and R square values for each association were normalized by the number of reads populating that contingency table. *python2 CoVaMa_Analyse_Matrices*.*py [Output_Tag]*.*Total_Matrices*.*py*.*pi CoVaMa_output*.*txt -- Min_Coverage 100 –-Min_Fusion_Coverage 100 -OutArray -Weighted NT*

### Statistical analysis in CoVaMa outcomes

Linkage disequilibrium was measured for every pair-wise interaction of each nucleotide position with each detected recombination event within the viral genome using the parameters set in the command line in each CoVaMa analysis (**Box1, Box2**). To distinguish the significant associations from the background noise, the three-sigma rule for comparing a single data point to a very large distribution of other data points was applied (27). CoVaMa automatically generated the standard deviation and mean of all LD values, based on which the three-sigma could be calculated. Associations passing the three-sigma in each CoVaMa outcome were considered significant.

## Results

### Description of algorithm

CoVaMa requires alignment information from standardized SAM files generated by any typical read mapper such as *bowtie2, hisat2* for short reads or *bwa* and *minimap2* for long reads. Specialized virus-focused read alignment software such as ViReMa can also be used if they output an alignment in SAM format. CoVaMa also requires information on the virus genome that was used in the alignment in FASTA format. The FASTA file provides information on the length of the viral genome and the number of genome segments. Using this information, CoVaMa builds three types of data matrices composed of numerous contingency tables. The largest is a Nucleotide-vs-Nucleotide matrix enumerating the pairwise nucleotide identities of every possible combination of mapped nucleotides found in every sequence read. The matrix comprises N*N potential contingency tables for each genome segment, where N is the length of the viral genome. Each contingency table is a 4×4 matrix where each row corresponds to the number of mapped A, T, G, or Cs in each read at the first genome coordinate, and each column corresponds to the number of mapped A, T, G, or Cs in each read at the second read coordinate, as we previously described (16).

In this manuscript, we introduced a novel feature in CoVaMa to correlate the presence or absence of recombination events including insertions, duplications, and deletions. We built matrices containing 4×2 Nucleotide-vs-Recombination contingency tables and 2×2 Recombination-vs-Recombination contingency tables. CoVaMa scrutinizes the provided SAM file for evidence of any recombination event (reported as an ‘N’, ‘D’ or ‘I’ in the CIGAR string) and adds the recombination event to the 4×2 and 2×2 matrices if it is mapped with at least a user-defined number of reads (10 by default). Similar to the Nucleotide-vs-Nucleotide tables, the columns of each 4×2 or 2×2 contingency table correspond to either the presence or the *confirmed* absence of a recombination event in each mapped read. These matrices and contingency tables are depicted in **Figure 1A**.

**Figure 1.**
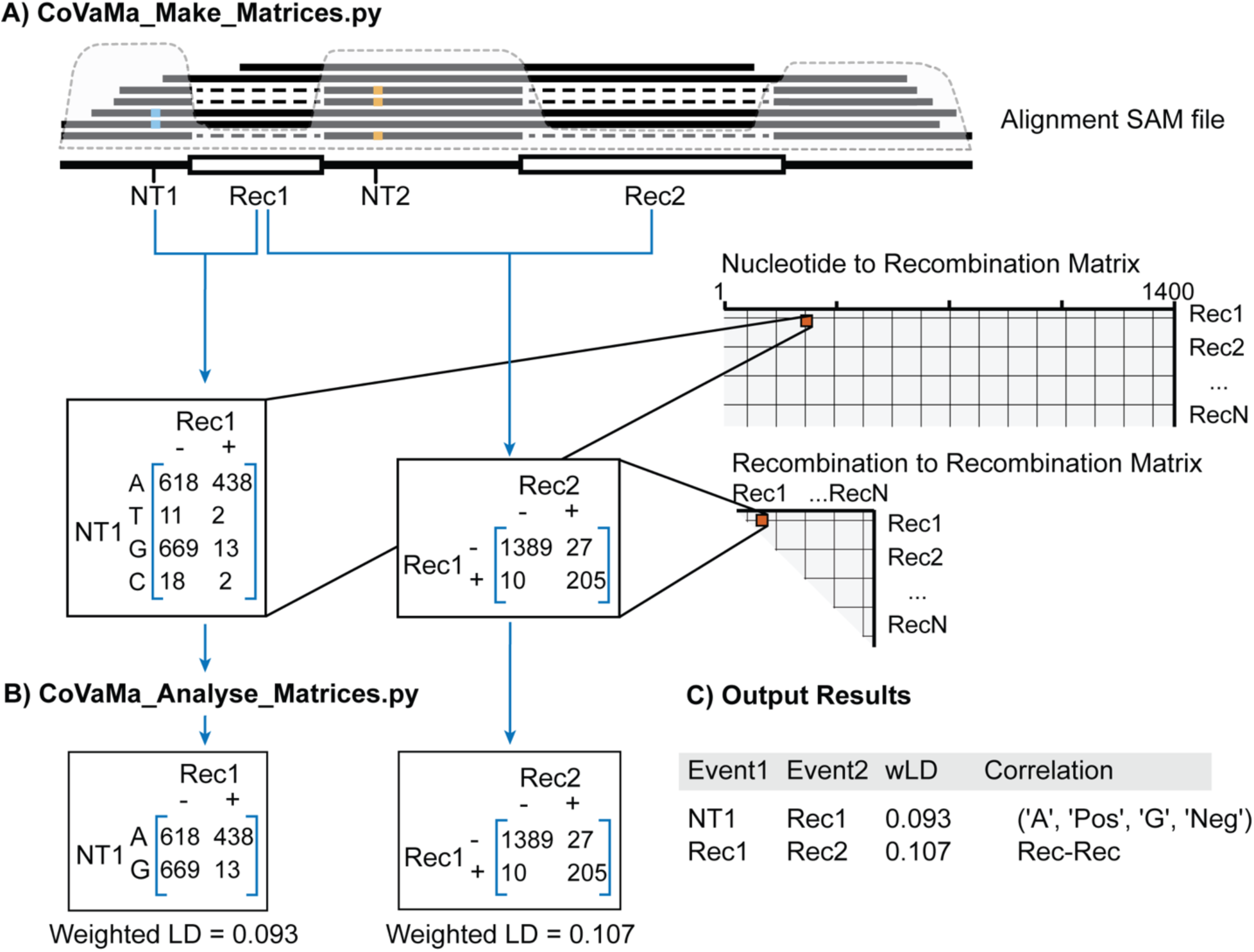
Schematic and flow chart of CoVaMa pipeline. **(A)** CoVaMa_Make_Matrices.py extracts information from each aligned read and generates large matrices containing 4×4 Nucleotide-vs-Nucleotide contingency tables, 4×2 Nucleotide-vs-Recombination contingency tables, and 2×2 Recombination-vs- Recombination contingency tables. In each contingency table, the columns correspond to either the mapped A, T, G, and Cs, for nucleotides, or the presence and the confirmed absence for recombination events. Rec, Recombination; SNV, single-nucleotide variant. **(B)** CoVaMa_Analyse_Matrices.py analyses contingency tables from each matrix for evidence of linkage disequilibrium. From each 4×4, 4×2, and 2×2 contingency table, every possible 2×2 table populated by sufficient reads is extracted to calculate the LD value. The LD values and R^2^ values are normalized by the proportion of reads populating the 2×2 contingency table. **(C)** An example of the CoVaMa output.

The contingency tables within each matrix are populated using all the individual reads from the provided SAM file using the ‘*CoVaMa_Make_Matrices*.*py’* python script. Linkage disequilibrium analyses rely on the measurement of both the presence of the associated alleles and their absence. For the 4×4 nucleotide matrices, the presence of one nucleotide at a locus automatically infers the absence of the other three. However, to measure linkage disequilibrium between recombination events, or between recombination events and nucleotides, we must build a pipeline that determines whether a mapped read can confirm that a recombination event is absent. To this end, we devised the following set of processes and parameters:

#### 1) Deletions/Splicing

To confirm the absence of a deletion event, a read must map inside the putative deletion site with at least a minimum number of nucleotides defined in the command-line (--Rec_Exclusion X; default is 5 nts). These mapped nucleotides can map anywhere within the deletion and/or overlap with the recombination junction. With a large number of nucleotides required, the exclusion of the recombination event has high confidence. If too few nucleotides are required, ambiguity may arise due to sequence similarity between deleted nucleotides and the nucleotides upstream of the recombination event. For instance, if only one nucleotide is required to map inside the recombination event, a negative mapping will be scored erroneously for one out of every four recombination events as the first deleted nucleotide has a one in four chance of also being the same as the nucleotide upstream of the recombination event. Care also must be taken in repetitive regions in case the nucleotides preceding the recombination 5’ site are close or identical to those preceding the recombination 3’ site. In cases like these, a large ‘X’ value is required.

#### 2) Micro-Deletions

If a deletion event is very small (i.e., it is a micro-deletion smaller than the --MicroInDel_Length Y parameter specified in the command line) then it is not possible for -- InDel_Exclusion X nts (default is 5) to map inside the recombination event. In this case, X nts are required to map continuously map on both sides of the recombination event to confirm the absence of the putative deletion.

#### 3) Insertions

A similar strategy is required to confirm the absence of a putative insertion event. Insertion events can either correspond to inserted nucleotides, insertions of fragments of host of other viral genes, or small duplications. To negate a large insertion event with length X, an aligned sequence should have at least X nucleotides on both sides of the possible insertion events. The length of required flanking sequences is controlled by the *Rec_Exclusion* value. An inappropriate *Rec_Exclusion* value decreases the number of negative events detected and might influence the analyzing results. Thus, insertion events are analyzed separately with different *Rec_Exclusion* values.

### Linkage Disequilibrium Calculation

Once contingency tables have been generated, evidence for covariation is determined using Linkage Disequilibrium (LD) by the same principle as we previously described, using the second python script called ‘*CoVaMa_Analyse_Matrices*.*py’*. From each 4×4, 4×2, and 2×2 contingency table, every possible 2×2 table is extracted. If the total number of reads for each 2×2 table exceeds a default value of x reads (which can be adjusted on the command-line), LD is calculated using the canonical formula: LD = (pAB * pab) – (pAb * paB) where ‘A’ and ‘a’ are the haplotypes for the nucleotide or recombination event at coordinate A, and ‘B’ and ‘b’ and the haplotypes for the nucleotide or recombination event at coordinate B, as depicted in **Figure 1B**. The range of LD is 0 to +/- 0.25, with higher values connoting disequilibrium and a value of 0 meaning there is no co-variation. In addition to the LD value, CoVaMa also calculates the R^2^ value using the canonical formula: R^2^ = (LD*LD)/ (pA * pa * pB * pb). The range of R^2^ is 0 to 1, again with higher values connoting disequilibrium and a value of 0 meaning no co-variation. CoVaMa also calculates the maximum possible LD that could have been obtained for each 2×2 contingency table given the frequencies of the variants at each coordinate. As a single 4×4 or 4×2 table can give rise to multiple possible LD reports if there is sufficient diversity in these contingency tables, as we described previously (16), the linkage disequilibrium values (LD) and R^2^ values for each association are normalized by the proportion of reads populating the 2×2 contingency table from the entire 4×4 or 4×2 contingency table. This yields weighted LD values (wLD) and weighted R^2^ values (wR^2^).

This information is reported in an output text file, as depicted in **Figure 1C**. This provides: 1) a description of the type of correlation being reported (e.g. nucleotide-vs-nucleotide, nucleotide-vs-recombination, or recombination-vs-recombination); 2) the two events being tested for correlation (either the nucleotide position or the recombination event); 3) the largest LD and R^2^value found the contingency table; 4) and largest possible LD values that could have been found given the frequencies of each observed variants at each coordinate; 5) and a flattened 2D-array of the contingency table that was tested for LD.

### Associations between recombination events and SNVs in defective FHV RNAs revealed by CoVaMa

Flock House virus (FHV) is an insect-specific small bipartite RNA virus and an ideal model system to study viral evolution and recombination (16, 28). The viral genome consists of two segments, RNA1 (3.1 kb) and RNA2 (1.4kb), which encode for the viral polymerase and viral capsid protein respectively (29). In a previous study (24), we serially passaged FHV in S2 Drosophila cells in culture to characterize the emergence, selection, and adaption of Defective-RNAs (D-RNAs) *in vitro*, which arise through nonhomologous RNA recombination (30). We began with a clonally derived population of viral RNAs expressed from transfected pMT vectors, and blind-passaged the supernatant every three days onto fresh S2 cells for a total of 9 passages. During this period, we extracted encapsidated RNA from purified virions and generated full-length cDNA copies of the FHV genomic segments using RT-PCR with primers designed for the first and last ∼25nts of each segment. Purified cDNA was prepared for long-read nanopore sequencing using the SQK-LSK107 2D kit to identify the emergence of insertions, deletions, and other RNA recombination events. We calculated an average error single-nucleotide mismatch rate of ∼7% in our 2D nanopore reads and we also robustly identified numerous recombination events including insertions and deletions. Interestingly, nanopore sequencing revealed that multiple deletions were most commonly found in individual reads, while reads containing only single-deletions were seldom seen. In that study, we postulated that the correlation of multiple deletion events within individual reads indicated a selective advantage for ‘mature’ D-RNAs over the ‘immature’ D-RNAs. The observation of these paired deletions was consistent with major defective RNA2 species that were previously characterized (24, 31–33).

During passaging, we also noticed the emergence of multiple minorities and SNVs in the viral genome including A226G and G575A in the RNA2. However, the function of these mutations was not determined. A226G is a synonymous substitution, while G575A results in the Alanine to Threonine substitution at amino acid position 185 of the capsid protein. This is at the five-fold and quasi-three-fold symmetry axes of the T=3 icosahedral virus particle (34) and might therefore interfere with the virus assembly (**SFig 1**).

To determine whether these SNVs co-varied with the recombination events constituting D-RNA2 species, we passed the mapped FHV nanopore reads from each passage to our CoVaMa pipeline using the command-line parameters indicated in **Box 1**. This used the aligned sequence data to populate contingency tables corresponding to the association of all mapped recombination events (found in at least 10 reads) with other recombination events or with SNVs (with a mutant frequency higher than 5%). As most erroneous InDels generated by Nanopore sequencing were found to be shorter than 25nts for this dataset (24), only recombination events greater than 25nts in length were used for analysis. We found that recombination events were infrequent in the first two passages, but multiple recombination events emerged in Passage 3 (**Fig 2A**). Throughout passaging, diverse recombination events were observed to increase in frequency, reflecting the selective or replicative advantage of Defective RNA species. Interestingly, the most common recombination events were clustered into two regions with only small variance in the exact coordinates of the recombination junction: deletion events that excise nucleotides between nt 240 and nt 530 termed ‘*Group 1*’ (such as 248^512 and 250^513), and deletion events that excised nucleotides between nt 720 and nt 1240 termed ‘*Group 2*’ (such as 736^1219) (**Fig 2B, Fig 2C**). The fluctuation in their abundance over time could be due to the competition between the encapsulation of D-RNAs and the requirement of generating enough complete capsid proteins to form mature capsids (35). As expected, neither group removed the RNA2 packing motif or the RNA2 replication *cis*-acting motif (32, 36).

**Figure 2:**
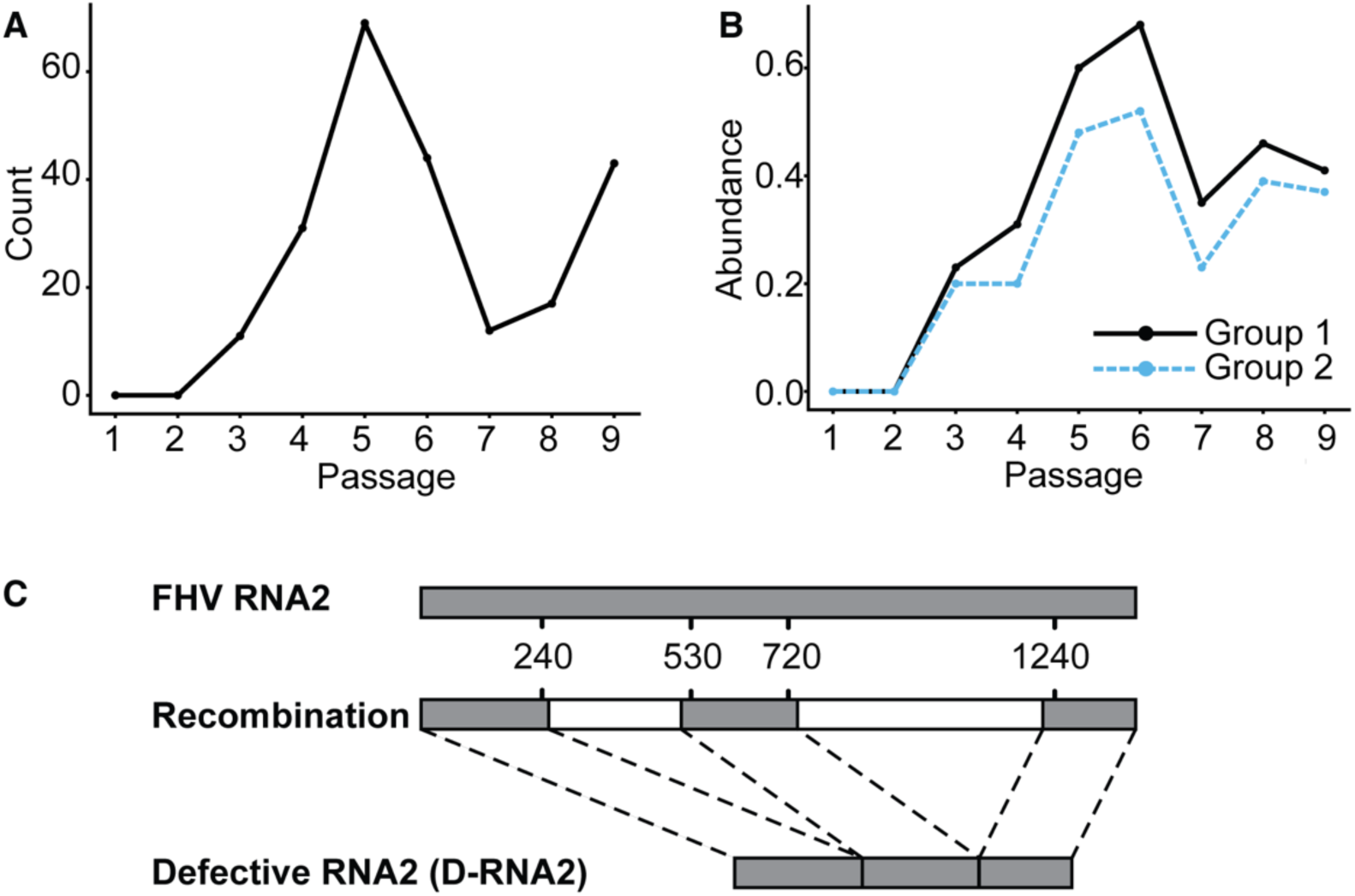
Recombination events detected in FHV RNA2 by CoVaMa. **(A)** The count of unique recombination events in FHV RNA2 in each passage. **(B)** The abundance of two groups of recombination events in the FHV RNA2 in each passage. **(C)** Schematic diagram of full-length FHV RNA2 and Defective RNA2 (D-RNA2).

CoVaMa detected strong associations between recombination events with each other and between recombination events and SNVs over passaging (**Fig 3A, Fig 3B, STable 1, SFig 2, STable 2**), including between 248^512 and 736^1219, 250^513 and 736^1219, G272A or A226G and recombination events. The non-synonymous mutation G575A positively correlated with recombination events 250^513, 248^512, and 736^1219 (**Fig 3B, SFig 2B**). This mutation was enriched over passaging along with the enrichment of D-RNA2, with a maintained significant LD between G575A and recombination events. This indicated that this SNV was predominantly found in the D-RNA2, but not full-length RNA2, suggesting a co-dependent evolution. In contrast to the G575A mutation, A226G mutation negatively correlated with recombination events 250^513, 248^512, and 736^1219 (**Fig 3B**). Consistent with these observations, CoVaMa also reported that A226G and G575A negatively correlated with one another (**Fig 3B, STable 3**).

**Figure 3:**
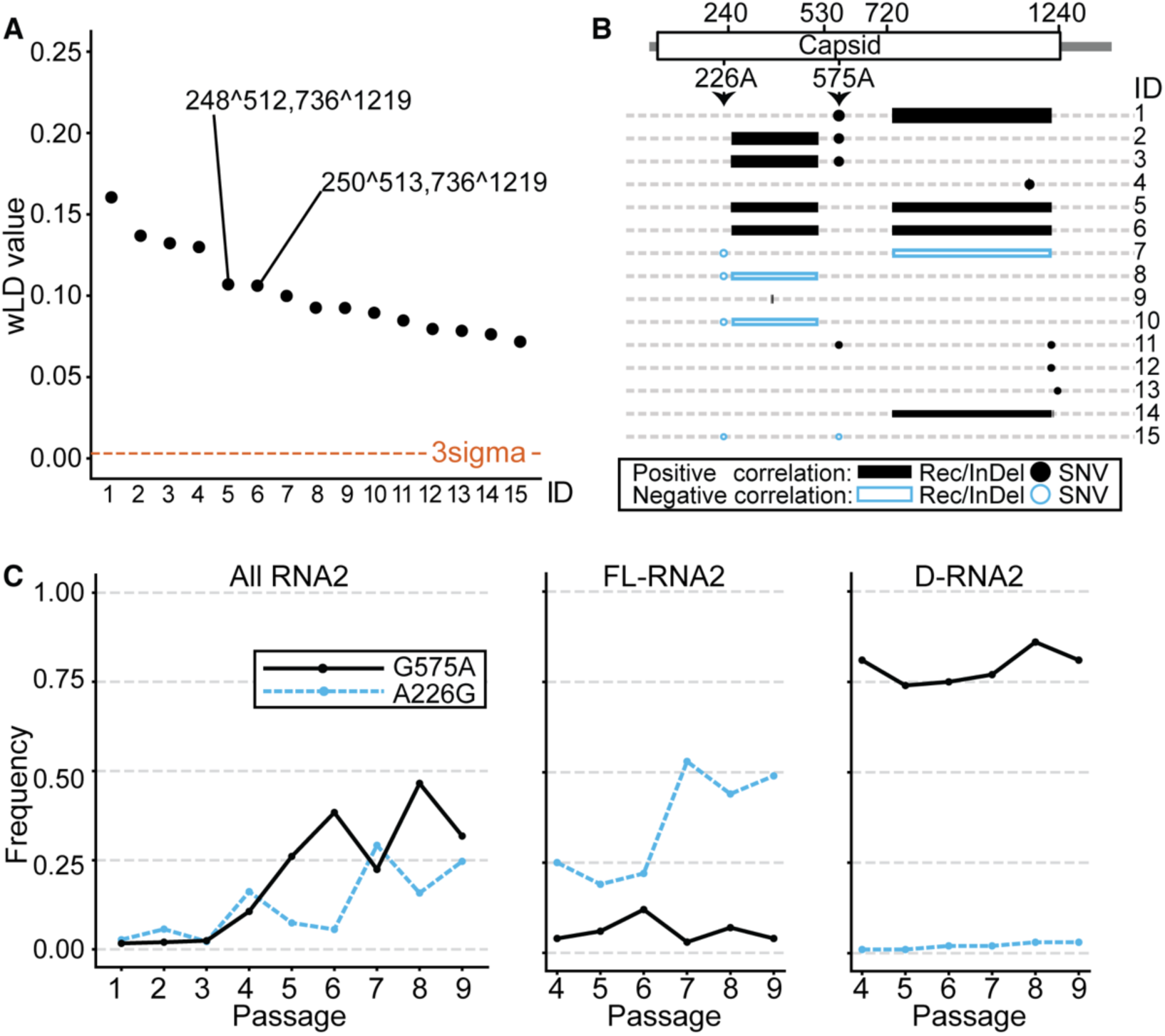
Associations between recombination events and SNVs in FHV RNA2 revealed by CoVaMa. The 15 associations with the highest wLD values in Passage 9 revealed by CoVaMa are labeled from 1 to 15. (**A**) The wLD values of the 15 associations are plotted from high to low. The associations between major recombination events are labeled in the plot. The three-sigma threshold is shown by the red dashed line. (**B**) This schematic diagram shows the distribution of the 15 associations on the FHV RNA2. Positive associations are colored in black while negative associations are colored in blue. Recombination events and InDels are plotted using blocks and SNVs are plotted using dots. The size of blocks and dots corresponds to the wLD value of each association. Recs, Recombination events. (**C**) The abundance of mutation A226G and G575A in all RNA2 reads (left), in the full-length RNA2 (middle), and in the D-RNA2 (right) in each passage.

To confirm these results, we separated reads mapping to either the full-length RNA2 or D-RNA2 into two groups based on whether they mapped over the recombination event 736^1219, which was one of the major recombination events constituting D-RNA2. By enumerating the frequency of A226G and G575A in full-length RNA2 and D-RNA2, we found that A226G was enriched in full-length RNAs from 25% in Passage 4 to 50% in Passage 9, while the abundance of A226G in D-RNA2 remained low at about 2% in all passages (**Fig 3C**). The ratio of the abundance of this mutation in full-length RNA2 and D-RNA2 was 18.8 ± 6.0 (SD) over passaging (**STable 4**). This SNV was therefore anti-correlated with the emergence of D-RNAs. The enrichment of A226G in the full-length RNA2 suggested a possible beneficial effect to the replication and/or packaging of the mutant full-length RNA2. In contrast, G575A was enriched in the D-RNA2 and relatively depleted in the full-length genomic RNA (**Fig 3C**). The ratio of the abundance of this mutation in D-RNA2 and full-length RNA2 was 16.2 ± 7.1 (SD) over passaging (**STable 5**). This SNV was therefore positively correlated with the emergence of D-RNAs, suggesting a D-RNA-specific adaptation.

A226G in RNA2 is a synonymous substitution, thus we investigated its potential influence on the secondary structure of RNA2. We used Vienna RNA Website (37) to generate predicted RNA structures of both D-RNA2 and full-length RNA2, with and without A226G. The presence of A226G showed no significant influence on the adjacent RNA2 packaging motif (32), both in D-RNA2 and full-length RNA2 (**SFig 3**). However, the A226G mutation was predicted to destabilize a region of secondary structure that formed a long-range interaction with residues 580-600, which in turn was predicted to alter additional long-range interactions even further downstream within the 50nts of the 3’ terminus. Interestingly, this region contains the cis-acting motif essential for RNA2 replication (38– 40). This might indicate a possible deleterious effect of A226G in the replication of D-RNA2. Alternatively, the predicted structure of full-length RNA2 without the A226G mutation placed position 736 and position 1219 close to each other, between which one of the major recombination events in D-RNA2 was found. However, when there was an A226G mutation in RNA2, this distance was significantly increased, which might preclude the emergence of this recombination event (**SFig 4**).

### Associations between insertion in p6^gag^ and mutations in *gag* cleavage sites revealed by CoVaMa

We reported on the covariation of amino acid and nucleotide variants within a large cohort of HIV-infected patients as part of the US Military HIV Natural History Longitudinal study using the previous version of CoVaMa (v0.1) (16). There, we found evidence of correlated mutations in the HIV *gag* and *protease* regions as well as multiple correlated adaptations within *protease* itself, similar to previous reports (6, 7, 16, 41, 42). However, these studies were limited to assessing only SNVs, and did not report on the presence of insertions, deletion, or duplication events, which are common in the HIV genome and critical for viral evolution (43). Here, we used ViReMa (23) to detect and quantify unusual or unknown insertions, deletion, and duplication events *de novo*. This approach leverages the seed-based mapping algorithm of *bowtie* and is particularly well suited to the analysis of complex viral RNA recombination events that fail to follow strict, or characterized, rules.

Five longitudinal sera samples were collected from a patient who continuously failed two antiretroviral treatments (ARTs) over six years (6). At each time point, HIV genomic RNA was RT-PCR amplified as previously described and sequenced using Next-Generation Sequencing (NGS) (26). The sequencing outputs were short reads 100nts long, which restricted the detection range of association to this length. We therefore merged paired reads using *BBMap*, extending the maximum detection range to 200 nucleotides. We analyzed the output NGS data from these longitudinal samples by mapping the data to the HXB2-indexed consensus genome of each sample and used ViReMa to identify recombination events in addition to SNVs that arose during virological failure. The average coverage over HIV *gag* and part of the *protease* was 40466 overlapping reads, with first and third quantile values being 23387 and 52143 reads. With this depth, we could identify numerous minority variants (**STable 6**) with at least 10 SNVs in each sample at a frequency of greater than 10%. In addition to SNVs, ViReMa reported a consistent insertion adding ‘RPEPS’ or ‘RLEPS’ in the P(T/S)AP region of the p6^gag^, which also generated a ‘RxEPS’ duplication (x for P/L) right downstream of the P1/p6 cleavage site (**Fig 4A, Table 1**). Similar insertion or duplication events in the p6 region close to the p1/p6 cleavage site have been observed in other studies (44–46). Another consistent 6-nt insertion occurred close to the proteolysis sites between matrix and capsid protein. This insertion encoded an extra ‘AA’ between amino acid 120 and amino acid 121 (**Fig 4A**), which has also been observed in the HIV 1 group M subgroup B isolate ARV2/SF2 (47). Additionally, a 15-nt insertion in the MA/CA cleavage site was detected in Sample 2 and Sample 4.

**Figure 4:**
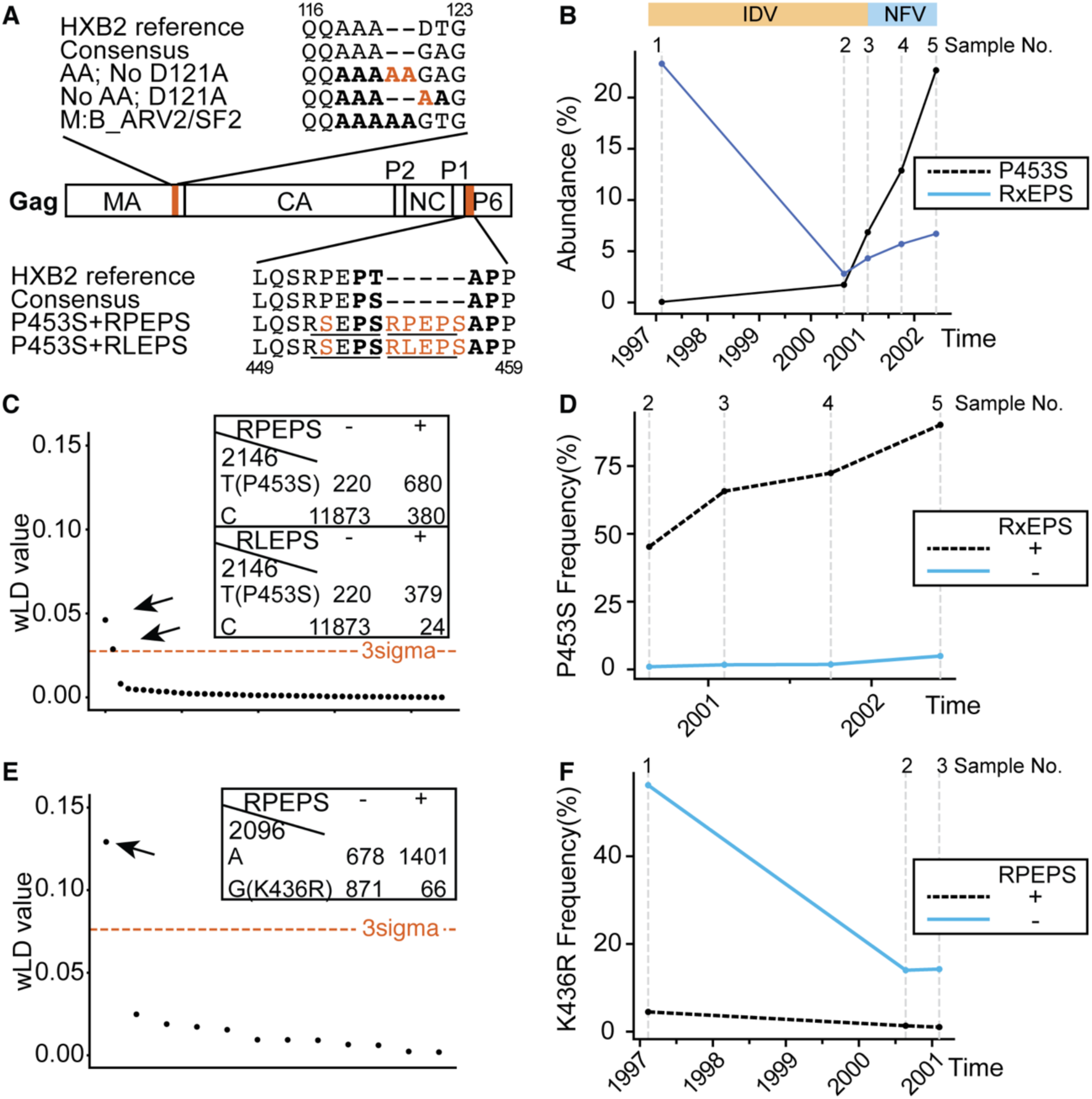
Associations between insertions in p6^Gag^ and mutations in Gag cleavage sites revealed by CoVaMa. **(A)** Positions of two major insertions detected and their correlated SNVs are shown in the HIV Gag, highlighted in red. Gag domains are indicated (MA, CA, P2, NC, P1, p6^Gag^). At the top are shown the HXB2 reference sequence, the consensus sequence of sample surrounding the ‘AA’ insertion between amino acid 120 and 121, the negative correlation between ‘AA’ and D121A, and the sequence for HIV 1 group M subgroup B isolate ARV2/SF2. The five-alanine motif is highlighted in bold. At the bottom are shown the HXB2 reference sequence, the consensus sequence of sample surrounding the ‘RxEPS’ insertion in p6^Gag^, and the positive correlations between ‘RxEPS’ and P453S (x for R/L). The PTAP region is highlighted in bold, and the duplication is marked underlined. **(B)** The abundance of ‘P453S’ and ‘RxEPS’ insertion in the viral population over time. The two antiretroviral therapies used were composed of Lamivudine (3TC), Zidovudine (AZT), and Indinavir (IDV) or Nelfinavir (NFV). **(C)** Significant correlations between ‘RxEPS’ insertions and P453S are indicated by arrows among all associations involving ‘RxEPS’ in Sample 4, with their contingency tables shown at the top right. **(D)** The abundance of P453S in reads with and without ‘RxEPS’ insertions. **(E)** Significant correlation between ‘RPEPS’ insertion and K436R is indicated by arrow among all associations involving ‘RPEPS’ in Sample 1, with its contingency table shown at the top right. **(F)** The abundance of K436R in reads with and without ‘RPEPS’ insertions.

**Table 1.**
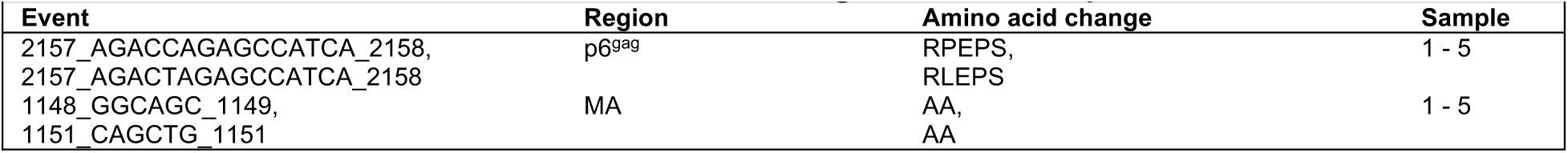
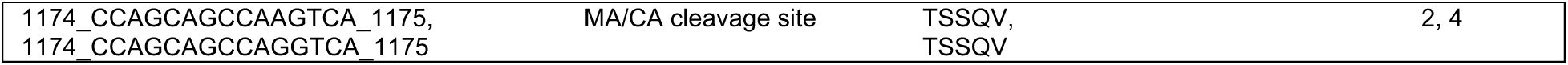
InDels and recombination events detected in longitudinal HIV samples.

The output SAM files from the ViReMa alignment were passed to CoVaMa to measure linkage disequilibrium. Significant associations were revealed between insertion events and SNVs. The C2146T mutation, which encodes the P453S substitution in the Gag P1/p6 cleavage site, emerged before sampling time 2, after which its abundance increased over time together with the enrichment of ‘RxEPS’ insertion in the p6^Gag^ (**Fig 4B**). CoVaMa showed that this mutation positively correlated with ‘RxEPS’ insertion (**Fig 4C, STable 7**), with a maintained elevated LD between them over time. Quantification of the abundance of P453S in reads with and without ‘RxEPS’ insertion showed that P453S was much more likely to be detected together with the ‘RxEPS’ insertion (**Fig 4D**). The ratio of P453S abundance in reads with and without ‘RLEPS’ was 27.74 ± 17.66 (SD) and the ratio of P453S abundance in reads with and without ‘RPEPS’ was 36.43 ± 16.67 (SD) (**STable 9**). In the last sample, while only 5% of the reads without ‘RxEPS’ insertion carried the P453S substitution, this number increased to 99% and 69% in reads with ‘RLEPS’ and ‘RPEPS’, respectively (**STable 9**). This finding supported cooperativity between P453S in the P1/p6 cleavage site and the ‘RxEPS’ insertion in the PTAP region. In contrast to C2146T, the A2096G mutation encoding the K436R substitution in the Gag NC/P1 cleavage site negatively correlated with the ‘RPEPS’ insertion (**Fig 4E, STable 8**). The ratio of the K436R abundance in reads without and with ‘RPEPS’ was 12.27 ± 1.71 (SD) in the first three samples before its abundance decreasing to lower than 1% (**Fig 4F, STable 9**). Consistent with the above, a combination of both K436R substitution and P453S substitution was not favored in the HIV genome (**STable 10**).

In addition to the associations between ‘RxEPS’ in p6^Gag^ and mutations in the Gag cleavages sites, the ‘AA’ insertion in the MA negatively correlated with the D121A substitution (**SFig 5A**), resulting in a five-alanine motif in the C terminus of MA, which has been seen in the isolate ARV2/SF2. Additionally, the ‘TSSQV’ insertion in the MA/CA cleavage site negatively correlated with D121A (**SFig 5B**) and positively correlated with the ‘AA’ insertion (wLD = 0.021805 in sample 4).

## Discussion

Adaption of viruses to their environments and to anti-viral therapies occurs through both the acquisition of novel single-nucleotide variants (SNVs) as well as recombination events such as small structural variants or larger insertions and deletions (18–20). These adaptions seldom occur in isolation, rather, multiple adaptions work together to confer the virus with novel biological properties. The covariation of SNVs during viral evolution has been well-described for a range of viral systems (6, 8). Bioinformatic tools that detect these co-varying SNVs are likewise readily available (reviewed in (9)). These report haplotypes using consensus-level data from large curated viral genomics databases as well as at the level of the viral intra-host diversity measured using Next-Generation Sequencing (13–15). However, computational tools that determine whether SNVs are correlated with recombination events or whether multiple recombination events are correlated with one another have to date been lacking. To address this gap, we therefore revised our previously reported CoVaMa (v0.1) pipeline to the new v0.7. We demonstrated the utility of this approach by studying two different viral systems characterized on different sequencing platforms.

Nanopore sequencing of FHV samples across passages revealed a strong linkage disequilibrium between the large deletion events that constituted D-RNA species that we previously characterized, consistent with the model that only ‘*mature’* D-RNAs efficiently replicate (24). Here we additionally found SNVs across the viral genome that were either positively correlated or negatively correlated with the deletions found in D-RNAs. These SNVs were previously too distantly spaced from each other or from deletion events to be correlated in the Illumina data, illustrating the value of performing long-read nanopore sequencing for this type of analysis.

The diversity and frequency changes of D-RNAs over passaging indicated the dynamic generation of D-RNAs from full-length RNAs and the competition among D-RNAs. Our data demonstrated that the D-RNA genomes acquired adaptions that were not found in the full-length wild-type genomic RNA. These adaptations in D-RNAs might be favored due to reduced genetic barriers of RNA2 mutation, due to not being responsible for functional viral expression, and could allow the formation of new secondary structures that confer D-RNAs with their-replicative and/or packaging advantage which have been granted the freedom to mutate by virtue of not being responsible for functional viral protein expression. Indeed, the G575A encodes an A185T mutation in the viral capsid protein at the five-fold and quasi-three-fold symmetry axes of the icosahedral virus particle. Such as bulky substitution at this interface may prevent efficient virus assembly. Conversely, adaptations in the full-length viral genome that were rarely found in the D-RNAs and only started to enrich after the emergence of D-RNAs (such as the synonymous A226G SNV found here), suggested that the full-length ‘*helper*’ virus may be adapting or escaping from the ‘*interfering’* properties of the D-RNAs. Although the mechanism of escape was not clear from these data, such a phenomenon was originally postulated by DePolo et al (48) who demonstrated that vesicular stomatitis virus (VSV) isolated from late viral passages was not subject to attenuation from defective interfering viral particles that arose in earlier viral passages. Further experimental characterization of these SNVs will reveal the precise molecular mechanisms driving their selection and competition.

Amino acid substitutions in cleavage sites and non-cleavage sites in HIV Gag have been shown to compensate for the compromised catalytic functions of protease with protease inhibitor (PI) Drug Resistant Mutations (DRMs) and contribute to PI sensitivity (6, 49–51). Besides the amino acid substitutions, insertions in Gag have been found to increase viral infectivity and drug resistance. Tamiya et al. reported that ‘SRPE’ duplication in p6^Gag^ in multi-PI resistant HIV subtype G could increase the cleavage efficiency of protease with DRMs (45). They also showed that several insertions, mostly duplications, near Gag p1/p6 cleavage site (e.g. APP duplication) in different multi-PI resistant HIV-1 subtypes could restore the compromised catalytic functions of mutant protease (45). Full or partial PTAP duplications have been reported to be selected during anti-retroviral treatment (52, 53). Martins et al. showed that, under PI pressure, full PTAP duplication in the Gag p6 significantly increases the cleavage efficiency of protease-bearing DRMs, thus increasing drug resistance and infectivity (54). In longitudinal serum samples collected from a patient who continuously failed antiretroviral therapies over six years, we detected an ‘R(P/L)EPS’ insertion in the P(T/S)AP region of the p6^Gag^. Using CoVaMa v0.7, we found this insertion in a strong correlation with mutations in Gag cleavage sites. The P453S mutation in the Gag P1/p6 CS was rarely detected in HIV strains without ‘R(P/L)EPS’ insertion, whereas it was enriched to over 90% in HIV isolates with ‘R(P/L)EPS’ insertion over time under drug pressure (**STable 9**). ‘RPEPS’ belongs to the proline-rich motif ‘RPEP(S/T)APP’ in the N terminus of p6^Gag^, which plays essential roles in the packaging of processed Pol proteins during late assembly (55, 56). Replacing the P453 and P455 showed reduced viral replication in primary monocytes (56), which might account for the consistent low frequency of P453S in viral strains we analyzed. However, the duplication of ‘RPEPS’ restores the two prolines or adds more prolines in this motif, which could release the restriction and allow the P453S substitution. Together, this might account for the strong positive correlation detected by CoVaMa between ‘RPEPS’ and P453S. Additionally, CoVaMa reported a weak correlation between the ‘RPEPS’ insertion in p6^Gag^ and A431V in the NC/P1 cleavage site (**STable 11**), which improves the cleavage efficacy and is mostly selected in the presence of protease DRM V82A (57, 58). The findings support the premise that for protease DRMs, duplications near the cleavage site and mutations in the cleavage site, may provide cooperative Gag cleavage functions and support drug-resistance development.

Overall, CoVaMa provides a simple and intuitive tool that probes both NGS datasets and nanopore datasets for evidence of the correlation between intra-host variants. Importantly, we here expanded this approach to detect and report the co-occurrence of SNVs with recombination events. While we focused here on viral intra-host diversity, the same approach and pipeline could equally be applied for NGS analysis of other organisms where diversity or correlation of sequence variants is anticipated, such as in bacterial or other complex mixtures of populations. We demonstrated the utility of this approach using both nanopore sequencing data acquired from the experimental evolution of the Flock House virus and using Illumina sequencing data acquired during the adaptation of HIV to antiviral therapies. In both cases, we observed novel associations of SNVs with specific recombination events. Knowledge of these associations is necessary to understand how viruses adapt to their environments and to characterize the distribution of specific genomic variants within viral intra-host diversity.

## Supporting information

Supplementary Data Part1

Supplementary Data Part2

## Data Availability

FHV datasets are publicly available in NCBI SRA under the accession code SRP094723. HIV datasets are publicly available in NCBI SRA under the accession code SRR15732332, SRR15732333, SRR15732334, SRR15732335, SRR15732336. The source code and supporting materials for CoVaMa (v0.7) are available at https://sourceforge.net/projects/covama/.

## Acknowledgments

We thank Dr. Yiyang Zhou for providing comments and critically reading the manuscript.

## Supplementary Data

Supplementary Data are available at NAR online.

