## Supplementary Data Part1 for "Co-variation of viral recombination with single nucleotide variants during virus evolution revealed by CoVaMa"

### Supplementary Table 1

| Supplementary Table 1. The 15 associations with the highest wLD values detected in FHV RNA2 in Passage 9. (3sigma = 0.002992) | | | | | | | |
|---|---|---|---|---|---|---|---|
| ID | NT1 | NT2 | wLD | wR2 | wLDmax | correlation | Contingency table |
| 1 | 575 | 736_to_1219 | 0.160452 | 0.649882 | 0.189137 | ('A', 'Pos', 'G', 'Neg') | [58, 478, 6, 7, 1364, 93, 7, 9] |
| 2 | 575 | 250_to_513 | 0.136847 | 0.648261 | 0.167558 | ('A', 'Pos', 'G', 'Neg') | [57, 342, 6, 7, 1367, 61, 7, 9] |
| 3 | 575 | 248_to_512 | 0.13225 | 0.593341 | 0.162764 | ('A', 'Pos', 'G', 'Neg') | [57, 338, 6, 2, 1367, 88, 7, 3] |
| 4 | 1152 | 1149_to_1152 | 0.129885 | 0.691951 | 0.134186 | ('T', 'Pos', 'C', 'Neg') | [6, 0, 23, 78, 0, 0, 356, 2] |
| 5 | 248_to_512 | 736_to_1219 | 0.106939 | 0.818994 | 0.123493 | Rec-Rec | [1389, 27, 10, 205] |
| 6 | 250_to_513 | 736_to_1219 | 0.106139 | 0.761762 | 0.122523 | Rec-Rec | [1389, 27, 24, 208] |
| 7 | 226 | 736_to_1219 | 0.099852 | 0.204686 | 0.110301 | ('A', 'Pos', 'G', 'Neg') | [640, 582, 12, 2, 640, 20, 16, 2] |
| 8 | 226 | 248_to_512 | 0.092589 | 0.190669 | 0.099929 | ('A', 'Pos', 'G', 'Neg') | [618, 438, 11, 2, 669, 13, 18, 2] |
| 9 | 370_to_372 | 374_to_376 | 0.092433 | 1.0 | 0.092433 | Rec-Rec | [235, 0, 0, 27] |
| 10 | 226 | 250_to_513 | 0.089469 | 0.184092 | 0.095802 | ('A', 'Pos', 'G', 'Neg') | [618, 406, 11, 2, 669, 11, 17, 3] |
| 11 | 575 | 1218 | 0.084728 | 0.426624 | 0.135626 | ('A', 'A', 'G', 'C') | [181, 8, 4, 85, 4, 0, 0, 6, 59, 12, 8, 1291, 4, 0, 0, 8] |
| 12 | 1217 | 1218 | 0.079553 | 0.794332 | 0.086432 | ('T', 'C', 'G', 'A') | [20, 4, 0, 13, 11, 2, 6, 1354, 143, 6, 2, 13, 2, 1, 0, 22] |
| 13 | 1237 | 1234_to_1237 | 0.07839 | 0.912376 | 0.080484 | ('T', 'Neg', 'C', 'Pos') | [4, 0, 1292, 3, 3, 0, 7, 124] |
| 14 | 736_to_1219 | 1223_to_1225 | 0.076266 | 0.164365 | 0.464003 | Rec-Rec | [486, 430, 0, 193] |
| 15 | 226 | 575 | 0.071722 | 0.125011 | 0.467635 | ('A', 'A', 'G', 'G') | [954, 20, 1143, 23, 11, 0, 11, 0, 38, 3, 650, 5, 4, 0, 23, 2] |

### Supplementary Table 2

| Supplementary Table 2. The 15 associations with the highest wLD values detected in FHV RNA2 in Passage 6 (3sigma = 0.006272) | | | | | | | |
|---|---|---|---|---|---|---|---|
| ID | NT1 | NT2 | wLD | wR2 | wLDmax | correlation | Contingency table |
| 1 | 250_to_513 | 736_to_1219 | 0.207513 | 0.709838 | 0.254205 | Rec-Rec | [134, 12, 8, 103] |
| 2 | 248_to_512 | 736_to_1219 | 0.20223 | 0.663083 | 0.246511 | Rec-Rec | [134, 12, 13, 112] |
| 3 | 1217 | 732_to_1216 | 0.169188 | 0.928924 | 0.174213 | ('T', 'Neg', 'G', 'Pos') | [3, 0, 150, 1, 1, 44, 0, 0] |
| 4 | 575 | 248_to_512 | 0.166334 | 0.47638 | 0.197724 | ('A', 'Pos', 'G', 'Neg') | [14, 208, 2, 3, 158, 53, 1, 7] |
| 5 | 575 | 250_to_513 | 0.163992 | 0.450745 | 0.197325 | ('A', 'Pos', 'G', 'Neg') | [14, 185, 2, 2, 158, 56, 1, 2] |
| 6 | 1217 | 1218 | 0.159645 | 0.705782 | 0.174138 | ('T', 'C', 'G', 'A') | [7, 1, 0, 6, 5, 6, 1, 216, 79, 5, 1, 9, 1, 0, 1, 7] |
| 7 | 1218 | 732_to_1216 | 0.159295 | 0.865189 | 0.164246 | ('A', 'Pos', 'C', 'Neg') | [3, 42, 3, 2, 0, 0, 151, 1] |
| 8 | 575 | 736_to_1219 | 0.147404 | 0.378178 | 0.193736 | ('A', 'Pos', 'G', 'Neg') | [24, 241, 1, 5, 168, 74, 3, 2] |
| 9 | 1218 | 250_to_513 | 0.134 | 0.474484 | 0.165746 | ('A', 'Pos', 'C', 'Neg') | [6, 39, 3, 1, 0, 4, 120, 16] |
| 10 | 249_to_517 | 730_to_1229 | 0.133563 | 0.373929 | 0.137253 | Rec-Rec | [143, 1, 58, 69] |
| 11 | 1218 | 248_to_512 | 0.128188 | 0.421029 | 0.159601 | ('A', 'Pos', 'C', 'Neg') | [6, 39, 3, 1, 0, 1, 120, 21] |
| 12 | 1218 | 734_to_1218 | 0.124059 | 0.80139 | 0.13453 | ('A', 'Pos', 'C', 'Neg') | [3, 29, 3, 1, 0, 0, 153, 2] |
| 13 | 736_to_1219 | 1223_to_1225 | 0.123355 | 0.340403 | 0.36238 | Rec-Rec | [48, 49, 0, 108] |
| 14 | 1217 | 248_to_512 | 0.115831 | 0.546442 | 0.193212 | ('T', 'Neg', 'G', 'Pos') | [4, 1, 121, 13, 1, 26, 0, 2] |
| 15 | 1224 | 734_to_1222 | 0.11202 | 0.432995 | 0.14202 | ('T', 'Neg', 'C', 'Pos') | [0, 0, 71, 3, 0, 1, 9, 16] |

### Supplementary Table 3

| Supplementary Table 5. A226G and G575A are negatively associated with each other in FHV RNA2. | | | | | | | |
|---|---|---|---|---|---|---|---|
| Passage | NT1 | NT2 | wLD | wR2 | wLDmax | correlation | Contingency table |
| 6 | 226 | 575 | 0.013454 | 0.014234 | 0.553561 | ('A', 'A', 'G', 'G') | [549, 16, 808, 13, 4, 0, 13, 0, 13, 1, 72, 1, 2, 0, 4, 0] |
| 9 | 226 | 575 | 0.071722 | 0.125011 | 0.467635 | ('A', 'A', 'G', 'G') | [954, 20, 1143, 23, 11, 0, 11, 0, 38, 3, 650, 5, 4, 0, 23, 2] |

### Supplementary Table 4

| Supplementary Table 4. Frequency of A226G in full-length RNA2 and D-RNA2 over passaging. | | |
|---|---|---|
| | A | G |

| Passage | All RNA2 | In full-length RNA2 | In D-RNA2 | All RNA2 | In full-length RNA2 | In D-RNA2 |
|---|---|---|---|---|---|---|
| 1 | 0.96 | NA | NA | 0.03 | NA | NA |
| 2 | 0.93 | NA | NA | 0.06 | NA | NA |
| 3 | 0.97 | NA | NA | 0.02 | NA | NA |
| 4 | 0.83 | 0.73 | 0.99 | 0.16 | 0.25 | 0.01 |
| 5 | 0.92 | 0.79 | 0.97 | 0.07 | 0.19 | 0.01 |
| 6 | 0.94 | 0.75 | 0.96 | 0.06 | 0.22 | 0.02 |
| 7 | 0.69 | 0.45 | 0.97 | 0.29 | 0.53 | 0.02 |
| 8 | 0.83 | 0.54 | 0.95 | 0.16 | 0.44 | 0.03 |
| 9 | 0.74 | 0.49 | 0.96 | 0.25 | 0.49 | 0.03 |

NOTE: The frequency of SNV in wild-type RNA2 and defective RNA2 is calculated from the contingency table formed between SNV and the major recombination event 736^1219, which is not available in Passage 1, 2, and 3 due to the low frequency of the SNV and/or the recombination event 736^1219. NA, not available.

### Supplementary Table 5

**Supplementary Table 3. Frequency of G575A in full-length RNA2 and D-RNA2 over passaging.**

| | A | | | G | | |
|---|---|---|---|---|---|---|
| Passage | All RNA2 | In full-length RNA2 | In D-RNA2 | All RNA2 | In full-length RNA2 | In D-RNA2 |
| 1 | 0.02 | NA | NA | 0.97 | NA | NA |
| 2 | 0.02 | NA | NA | 0.97 | NA | NA |
| 3 | 0.02 | NA | NA | 0.97 | NA | NA |
| 4 | 0.11 | 0.04 | 0.81 | 0.89 | 0.95 | 0.17 |
| 5 | 0.26 | 0.06 | 0.74 | 0.73 | 0.93 | 0.24 |
| 6 | 0.38 | 0.12 | 0.75 | 0.61 | 0.86 | 0.23 |
| 7 | 0.22 | 0.03 | 0.77 | 0.77 | 0.95 | 0.17 |
| 8 | 0.47 | 0.07 | 0.86 | 0.52 | 0.90 | 0.11 |
| 9 | 0.32 | 0.04 | 0.81 | 0.68 | 0.95 | 0.16 |

NOTE: The frequency of SNV in wild-type RNA2 and defective RNA2 is calculated from the contingency table formed between SNV and the major recombination event 736^1219, which is not available in Passage 1, 2, and 3 due to the low frequency of the SNV and/or the recombination event 736^1219.  NA, not available.

### Supplementary Table 6

**Supplementary Table 6. Common SNVs in each sample.**

| Locus | Ref | Alt | Alt_Frequency | Locus | Ref | Alt | Alt_Frequency | Locus | Ref | Alt | Alt_Frequency |
|---|---|---|---|---|---|---|---|---|---|---|---|
| **Sample No.1: Feb 1997** | | | | **Sample No.2: Aug 2000** | | | | **Sample No.3: Feb 2001** | | | |
| 888 | T | C | 0.4 | 1033 | G | A | 0.57 | 832 | C | A | 0.51 |
| 915 | A | G | 0.3 | 909 | G | A | 0.44 | 1033 | G | A | 0.49 |
| 2442 | A | G | 0.26 | 1151 | A | G | 0.43 | 915 | A | G | 0.47 |
| 1995 | C | T | 0.23 | 915 | A | G | 0.4 | 1030 | A | G | 0.31 |
| 2466 | G | A | 0.23 | 960 | T | C | 0.29 | 1717 | T | A | 0.3 |
| 2436 | G | A | 0.16 | 834 | A | G | 0.28 | 1809 | G | A | 0.23 |
| 2497 | C | T | 0.16 | 852 | A | G | 0.27 | 960 | T | C | 0.19 |
| 1160 | A | C | 0.14 | 1809 | G | A | 0.26 | 909 | A | G | 0.16 |
| 1881 | G | A | 0.13 | 1030 | A | G | 0.22 | 888 | T | C | 0.12 |
| 2221 | A | C | 0.08 | 1717 | T | A | 0.17 | 1995 | C | T | 0.11 |
| **Sample No.4: Oct 2001** | | | | **Sample No.5: Jun 2002** | | | | | | | |
| 1717 | T | A | 0.54 | 2390 | G | A | 0.42 | | | | |
| 1030 | A | G | 0.39 | 1263 | A | C | 0.41 | | | | |
| 1033 | G | A | 0.39 | 1977 | C | T | 0.41 | | | | |
| 1155 | A | C | 0.36 | 889 | A | G | 0.37 | | | | |
| 832 | C | A | 0.33 | 2467 | T | C | 0.34 | | | | |
| 2412 | A | G | 0.33 | 1432 | G | T | 0.31 | | | | |
| 2517 | C | A | 0.2 | 1479 | A | G | 0.29 | | | | |
| 2534 | T | C | 0.19 | 978 | G | A | 0.29 | | | | |
| 1263 | A | C | 0.18 | 1356 | C | T | 0.28 | | | | |
| 1809 | G | A | 0.17 | 1995 | C | T | 0.26 | | | | |

NOTE: NGS data from each sample is mapped to the HXB2-indexed consensus genome of that sample.

### Supplementary Table 7

**Supplementary Table7. Associations involving the RxEPS insertion in the p6$^{Gag}$ detected by CoVaMa in Sample 4. (3sigma = 0.027592)**

| NT1 | NT2 | wLD | wR2 | wLDmax | correlation | Contingency table |
|---|---|---|---|---|---|---|

| | | | | | | |
|---|---|---|---|---|---|---|
| 2146 | 2157_AGACCAGAGCCATCA_2158 | 0.046153 | 0.451311 | 0.062868 | ('T', 'Pos', 'C', 'Neg') | [3, 0, 220, 680, 6, 0, 11873, 380] |
| 2146 | 2157_AGACTAGAGCCATCA_2158 | 0.028763 | 0.581251 | 0.030682 | ('T', 'Pos', 'C', 'Neg') | [3, 0, 220, 379, 6, 0, 11873, 24] |
| 2211 | 2157_AGACCAGAGCCATCA_2158 | 0.00815 | 0.007943 | 0.05387 | ('A', 'Pos', 'G', 'Neg') | [1364, 175, 0, 0, 7310, 424, 1, 0] |
| 2201 | 2157_AGACCAGAGCCATCA_2158 | 0.005183 | 0.006018 | 0.005183 | ('T', 'Neg', 'C', 'Pos') | [0, 0, 722, 0, 2, 0, 8702, 742] |
| 2198 | 2157_AGACCAGAGCCATCA_2158 | 0.004705 | 0.006266 | 0.053864 | ('T', 'Pos', 'C', 'Neg') | [1, 0, 529, 94, 1, 0, 9481, 655] |
| 2200 | 2157_AGACCAGAGCCATCA_2158 | 0.0045 | 0.004878 | 0.004792 | ('A', 'Pos', 'G', 'Neg') | [8836, 746, 0, 0, 670, 3, 1, 1] |
| 2100 | 2157_AGACCAGAGCCATCA_2158 | 0.004057 | 0.004686 | 0.004057 | ('T', 'Neg', 'C', 'Pos') | [0, 0, 123, 0, 0, 0, 2502, 278] |
| 2197 | 2157_AGACTAGAGCCATCA_2158 | 0.003539 | 0.01154 | 0.027437 | ('A', 'Neg', 'G', 'Pos') | [9742, 249, 3, 0, 375, 49, 1, 0] |
| 2214 | 2157_AGACCAGAGCCATCA_2158 | 0.003504 | 0.010557 | 0.018263 | ('A', 'Pos', 'G', 'Neg') | [133, 43, 1, 0, 8291, 542, 1, 0] |
| 2211 | 2157_AGACTAGAGCCATCA_2158 | 0.002959 | 0.002832 | 0.003747 | ('A', 'Neg', 'G', 'Pos') | [1364, 7, 0, 0, 7310, 209, 1, 0] |
| 2206 | 2157_AGACCAGAGCCATCA_2158 | 0.002584 | 0.002428 | 0.003092 | ('T', 'Pos', 'C', 'Neg') | [3, 0, 8728, 662, 1, 0, 444, 5] |
| 2147 | 2158_AGACCAGAGCCATCA_2158 | 0.002247 | 0.002515 | 0.002247 | ('T', 'Neg', 'C', 'Pos') | [3, 0, 367, 0, 8, 0, 11724, 1060] |
| 2169 | 2157_AGACCAGAGCCATCA_2158 | 0.002185 | 0.003019 | 0.080219 | ('A', 'Neg', 'G', 'Pos') | [11406, 989, 3, 0, 222, 50, 3, 1] |
| 2201 | 2157_AGACTAGAGCCATCA_2158 | 0.002117 | 0.002355 | 0.002117 | ('T', 'Neg', 'C', 'Pos') | [0, 0, 722, 0, 2, 0, 8702, 276] |
| 2200 | 2157_AGACTAGAGCCATCA_2158 | 0.001959 | 0.002165 | 0.001959 | ('A', 'Pos', 'G', 'Neg') | [8836, 280, 0, 0, 670, 0, 1, 0] |

### Supplementary Table 8

**Supplementary Table8. Associations involving the RxEPS insertion in the p6$^{Gag}$ detected by CoVaMa in Sample 1. (3sigma = 0.076140)**

| NT1 | NT2 | wLD | wR2 | wLDmax | correlation | Contingency table |
|---|---|---|---|---|---|---|
| 2096 | 2157_AGACCAGAGCCATCA_2158 | 0.129189 | 0.312063 | 0.151065 | ('A', 'Pos', 'G', 'Neg') | [678, 1401, 0, 0, 871, 66, 0, 1] |
| 2081 | 2157_AGACCAGAGCCATCA_2158 | 0.024842 | 0.038874 | 0.03672 | ('T', 'Pos', 'C', 'Neg') | [0, 0, 26, 124, 0, 0, 1147, 892] |
| 2221 | 2157_AGACCAGAGCCATCA_2158 | 0.018924 | 0.020563 | 0.026611 | ('A', 'Pos', 'C', 'Neg') | [4448, 1873, 4, 1, 16, 3, 625, 54] |
| 2166 | 2157_AGACCAGAGCCATCA_2158 | 0.017251 | 0.02066 | 0.349795 | ('A', 'Neg', 'G', 'Pos') | [7116, 3936, 2, 3, 283, 496, 0, 0] |
| 2097 | 2157_AGACCAGAGCCATCA_2158 | 0.015503 | 0.028963 | 0.016443 | ('A', 'Pos', 'G', 'Neg') | [1562, 1521, 0, 0, 107, 3, 0, 0] |
| 2155 | 2157_AGACCAGAGCCATCA_2158 | 0.00944 | 0.015457 | 0.00944 | ('A', 'Neg', 'T', 'Pos') | [301, 0, 7141, 4435, 0, 0, 30, 0] |
| 2130 | 2157_AGACCAGAGCCATCA_2158 | 0.009365 | 0.019293 | 0.00949 | ('T', 'Neg', 'C', 'Pos') | [0, 0, 148, 1, 0, 0, 3775, 4113] |
| 2136 | 2157_AGACCAGAGCCATCA_2158 | 0.009121 | 0.01863 | 0.009236 | ('T', 'Neg', 'C', 'Pos') | [0, 1, 158, 1, 0, 1, 4144, 4432] |
| 2148 | 2157_AGACCAGAGCCATCA_2158 | 0.006533 | 0.008644 | 0.008044 | ('A', 'Pos', 'C', 'Neg') | [7213, 4410, 1, 2, 20, 5, 239, 18] |
| 2190 | 2157_AGACCAGAGCCATCA_2158 | 0.006102 | 0.008152 | 0.007077 | ('A', 'Neg', 'G', 'Pos') | [203, 10, 0, 2, 6558, 3487, 0, 1] |
| 2211 | 2157_AGACCAGAGCCATCA_2158 | 0.002351 | 0.001924 | 0.009928 | ('A', 'Pos', 'G', 'Neg') | [62, 53, 0, 0, 5718, 2349, 1, 0] |
| 2100 | 2157_AGACCAGAGCCATCA_2158 | 0.001952 | 0.000289 | 0.029112 | ('T', 'Pos', 'C', 'Neg') | [0, 0, 94, 100, 0, 1, 1703, 1563] |

### Supplementary Table 9

**Frequency of SNVs in reads with and without 'RLEPS' insertion in longitudinal samples.**

| | Frequency of A2096G (K436R) in the NC/P1 cleavage site (%) | | | Frequency of C2146T (P453S) in the p1/p6 cleavage site (%) | | |
|---|---|---|---|---|---|---|
| Sample No. | In all reads | In reads without insertion | In reads with insertion | In all reads | In reads without insertion | In reads with insertion |
| 1 | 26 | NA | NA | 0 | NA | NA |
| 2 | 9 | 14.01 | 6.56 | 2 | 0.97 | 27.95 |
| 3 | 10 | 14.26 | 5.41 | 7 | 1.69 | 17.42 |

| | | | | | | |
|---|---|---|---|---|---|---|
| 4 | 1 | NA | NA | 13 | 1.82 | 94.04 |
| 5 | 0 | NA | NA | 23 | 4.92 | 99.18 |
| **Frequency of SNVs in reads with and without 'RLPEPS' insertion in longitudinal samples** | | | | | | |
| | Frequency of A2096G (K436R) in the NC/P1 cleavage site (%) | | | Frequency of C2146T (P453S) in the p1/p6 cleavage site (%) | | |
| Sample | In all reads | In reads without insertion | In reads with insertion | In all reads | In reads without insertion | In reads with insertion |
| 1 | 26 | 56.23 | 4.50 | 0 | NA | NA |
| 2 | 9 | 14.01 | 1.34 | 2 | 0.97 | 51.61 |
| 3 | 10 | 14.26 | 1.03 | 7 | 1.69 | 73.15 |
| 4 | 1 | NA | NA | 13 | 1.82 | 64.15 |
| 5 | 0 | NA | NA | 23 | 4.92 | 68.84 |

NOTE: The frequency of SNVs in reads with and without 'RxEPS' insertion is calculated from the contingency table, which is not available when the frequency of the SNV and/or insertion event is lower than the detection level. NA, not available.

### Supplementary Table 10.

**Supplementary Table 10. Contingency table for the association between 2096 and 2146 (sample 3).**

| 2096 \ 2146 | T (P453S) | C |
|---|---|---|
| A | 439 | 3302 |
| G (K436R) | 14 | 590 |

### Supplementary Table 11

**Supplementary Table 12. Contingency table for the association between C2081T and 'RPEPS' insertion (sample 1).**

| 2081 \ 'RPEPS' insertion | Negative | Positive |
|---|---|---|
| T (A431V) | 26 | 124 |
| C | 1147 | 892 |

**Supplementary Figure 1. A185T in the capsid protein locates at the quasi-three-fold and five-fold symmetry axes of the Flock House virus particle**. This capsid structure depicts the location of G575A (Alanine to Threonine substitution at amino acid 185) at quasi-three-fold and five-fold symmetry axes of the Flock House virus particle. The structure is modified based on 4FTB.

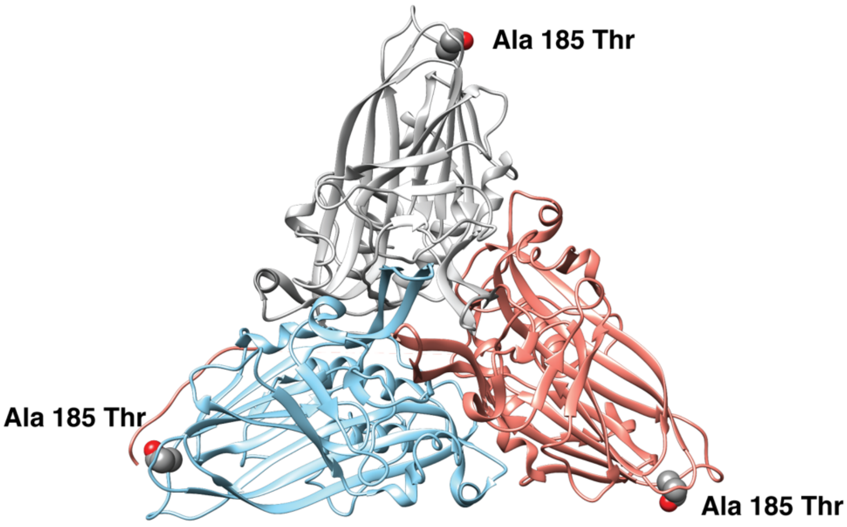

**Supplementary Figure 2. Associations between recombination events and SNVs in FHV RNA2 in Passage 6.** The 15 associations with the highest wLD values in Passage 6 revealed by CoVaMa are labeled from 1 to 15. **(A)** The wLD values of the 15 associations are plotted from high to low. The associations between major recombination events are labeled in the plot. The three-sigma threshold is shown by the red dashed line. **(B**) This schematic diagram shows the distribution of the 15 associations on the FHV RNA2. Recombination events and InDels are plotted using blocks and SNVs are plotted using dots. The size of blocks and dots corresponds to the wLD value of each association. Recs, Recombination events.

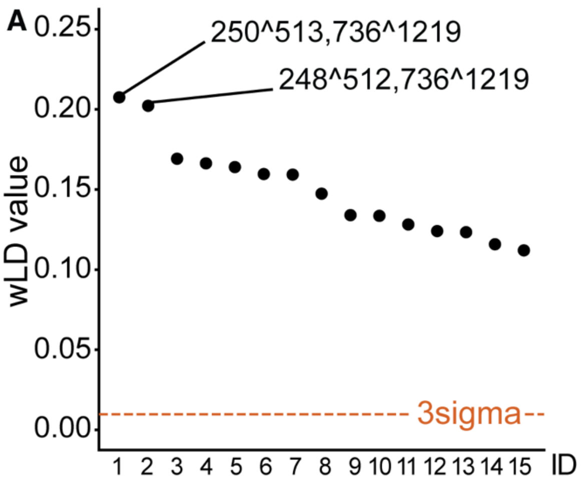
