## Supplementary Data Part2 for "Co-variation of viral recombination with single nucleotide variants during virus evolution revealed by CoVaMa"

**Supplementary Figure 3. Predicted FHV RNA2 structure with and without A226G.** From top to bottom are the predicted RNA structures of full-length RNA2 without A226G, full-length RNA2 with A226G, D-RNA2 without A226G, and D-RNA2 with A226G. The secondary structures are predicted using Vienna RNA Websuite and viewed in Forna (Gruber et al., 2008). The nucleotide at nt 226 is highlighted in red. The packing signal region at the 5' terminus is colored in blue, and the *cis-acting* signal at the 3' terminus is colored in orange. The predicted long-range interaction between nt 590-600 and nt 218-241 is colored in grey.

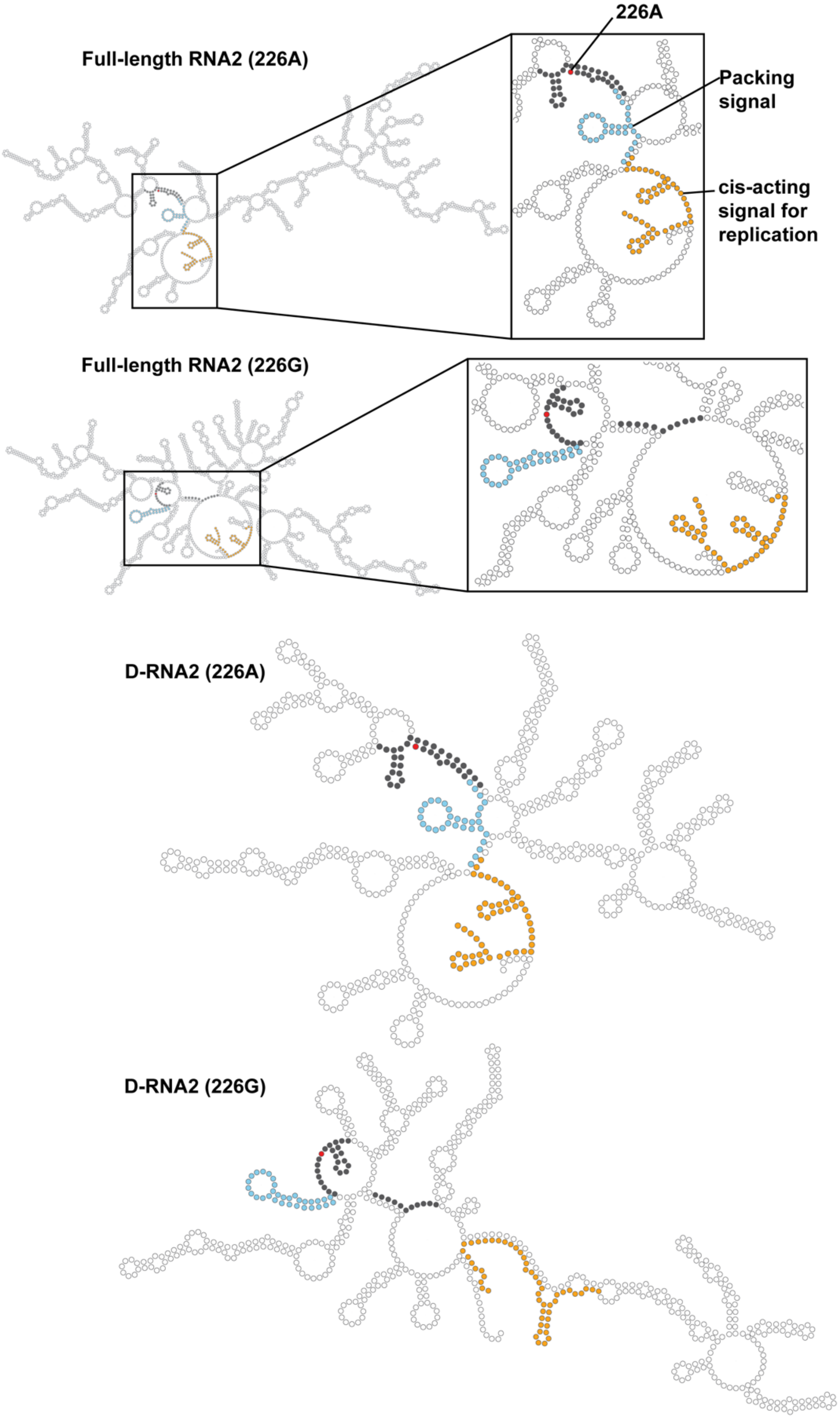
Full-length RNA2 (226A)
226A
Packing signal
cis-acting signal for replication
Full-length RNA2 (226G)
D-RNA2 (226A)
D-RNA2 (226G)

**Supplementary Figure 4. The recombination sites in predicted FHV RNA2 structure with and without A226G.** From top to bottom are the predicted RNA structures of full-length RNA2 without A226G and with A226G. The secondary structures are predicted using Vienna RNA Websuite and viewed in Forna (Gruber et al., 2008). Nucleotides at nt 736 and nt 1219 are highlighted in red.

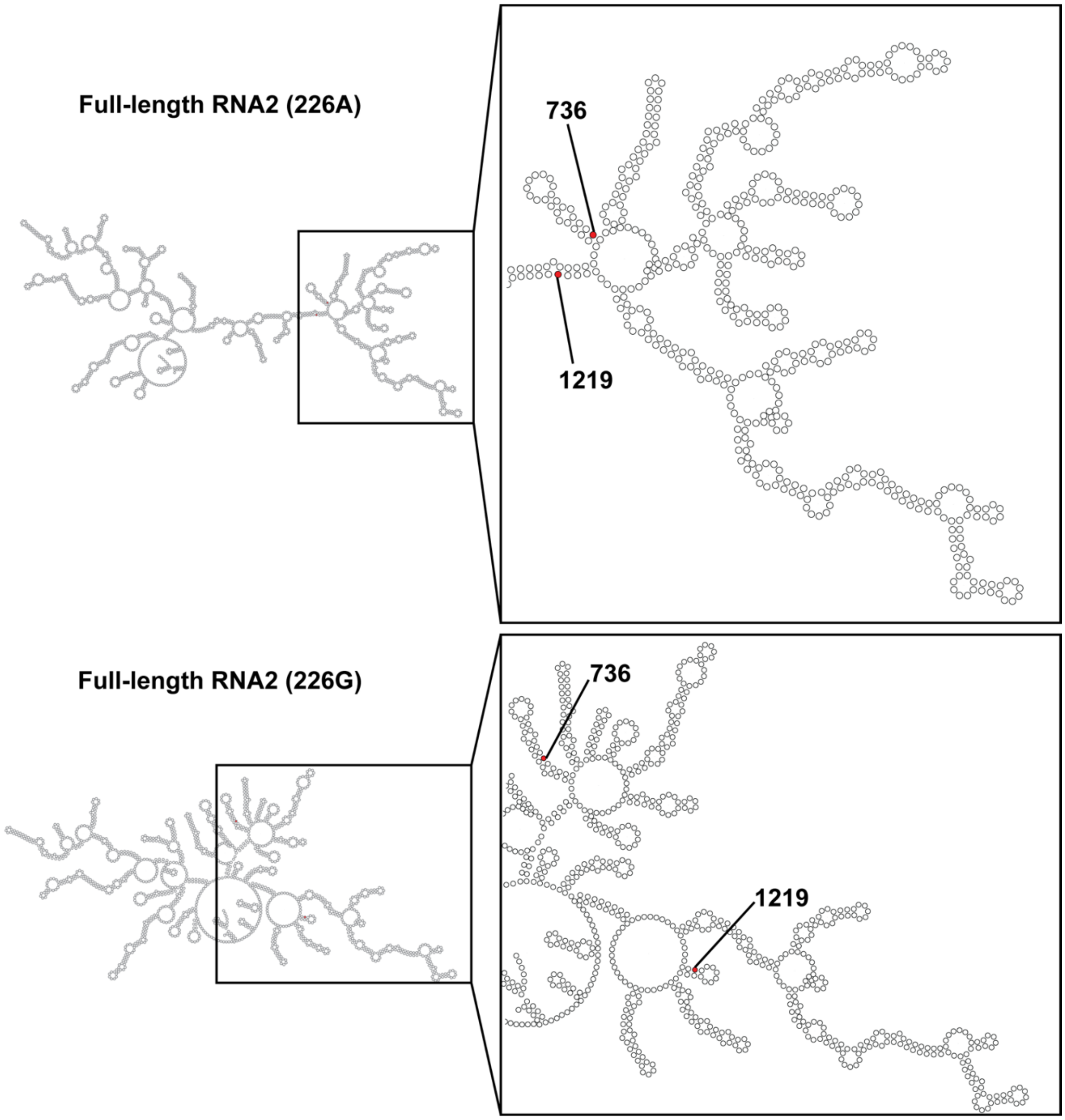

**Supplementary Figure 5. Associations between MA insertions and SNVs in the HIV genome. (A)** Associations involving the ‘AA’ insertion in the C-terminus of MA in Sample 3 are revealed by CoVaMa. The wLD values of the associations are plotted from high to low. The contingency table between this insertion and nt 1151 is shown on the top right. (B) Associations involving the ‘TSSQV’ insertion in the MA/CA cleavage site in Sample 4 are revealed by CoVaMa. The wLD values of the associations are plotted from high to low. The contingency table between this insertion and nt 1151 is shown on the top right. The three-sigma threshold is shown by the red dashed line.

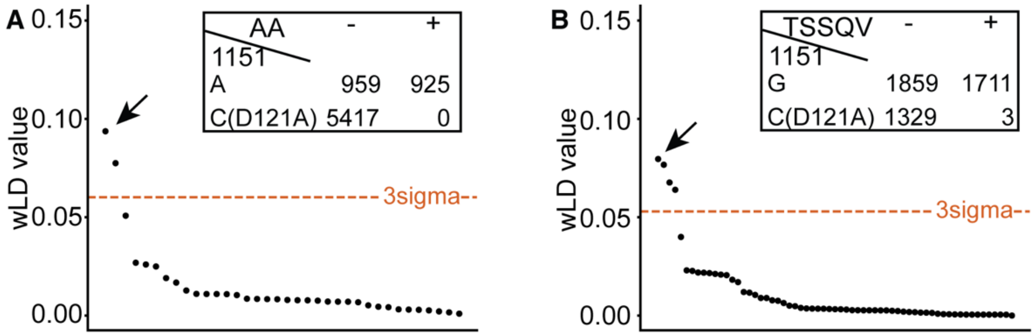
